# fCUT&Tag-Seq: An Optimized CUT&Tag-based Method for High-Resolution Profiling of Histone Modifications and Chromatin-Binding Proteins in Fungi

**DOI:** 10.1101/2025.01.23.634238

**Authors:** Haiting Wang, Yongjunlin Tan, Jiayue Ma, Jie Yang, Mengran Liu, Peng Jiang, Shan Lu, Haoxue Xia, Guangfei Tang, Wende Liu, Hui-Shan Guo, Chun-Min Shan

**Affiliations:** Department of Agri-microbiomics and Biotechnology, State Key Laboratory of Microbial Diversity and Innovative Utilization, Institute of Microbiology, Chinese Academy of Sciences, Beijing 100101, China; Shanxi Agricultural University, Jinzhong 030801, China; University of Chinese Academy of Sciences, Beijing 100049, China; College of Agriculture, Guangxi Key Laboratory of Sugarcane Biology, Guangxi University, Nanning 530004, China; State Key Laboratory for Biology of Plant Diseases and Insect Pests, Institute of Plant Protection, Chinese Academy of Agricultural Sciences, Beijing 100193, China

**Keywords:** fCUT&Tag-Seq, Fungi, Histone modifications, Chromatin-binding proteins, Plant pathogen

## Abstract

Histone modifications and chromatin-binding proteins play crucial roles in regulating gene expression in eukaryotes, with significant implications for fungal pathogenicity and development. However, profiling these modifications or proteins across the genome in fungi remains challenging due to the technical limitations of the traditional, widely used ChIP-Seq method. Here, we present an optimized CUT&Tag-Seq protocol (fCUT&Tag-Seq) specifically designed for filamentous fungi and dimorphic fungi. Our approach involves the preparation of protoplasts and nuclear extraction to enhance antibody accessibility, along with formaldehyde crosslinking to improve protein-DNA binding efficiency. We then successfully applied fCUT&Tag-Seq to accurately profile multiple histone modifications like H3K9me3, H3K27me3, H3K4me3, and H3K18ac, across different plant pathogenic or model fungal species, including *Verticillium dahliae*, *Neurospora crassa*, *Fusarium graminearum*, and *Sporisorium scitamineum*, showing good signal-to-noise ratios, reproducibility, and detection sensitivity. Furthermore, we extended this method to profile chromatin-binding proteins, such as the histone acetyltransferase Gcn5. This study establishes fCUT&Tag-Seq as a robust and useful tool for fungal epigenetic research, enabling detailed exploration of chromatin dynamics and advancing our understanding of fungal gene regulation, development, and pathogenicity.

**Impact Statement:** We developed a faster, lowLJinput method to study how genes are turned on and off in fungi, even in tough species that are difficult to analyze with standard methods. Our new approach, named fCUT&Tag-Seq, requires only 10,000 cells and can be completed in just two days. It delivers clearer, more reliable results and has already been successfully applied to multiple fungi, including crop pathogens. By revealing the molecular “control switches” that govern fungal development and virulence, we expect it will accelerate basic research and help identify new targets for controlling destructive plant diseases.

## Introduction

Chromatin structure and modifications are essential in regulating gene expression in eukaryotic organisms. The basic unit of chromatin, the nucleosome, consists of DNA wrapped around histone octamers composed of core histones H2A, H2B, H3, and H4[1]. The N-terminal tails of these histones can undergo various post-translational modifications (PTMs), such as methylation, acetylation, phosphorylation, ubiquitination, sumoylation, and ADP-ribosylation[2, 3]. These modifications can alter chromatin structure and impact vital biological processes, such as gene expression, DNA repair, and chromosome organization[4, 5].

Among these modifications, histone methylation is critical in regulating gene expression. For instance, trimethylation of histone H3 at lysine 9 (H3K9me3) and lysine 27 (H3K27me3) is highly associated with heterochromatin formation and gene silencing. In contrast, trimethylation of histone H3 at lysine 4 (H3K4me3), which typically occurs at transcription initiation sites, is commonly enriched near gene promoters and is associated with active transcription[6–8]. Furthermore, chromatin-binding proteins, such as histone acetyltransferases and transcription factors, play crucial roles in modifying histones or binding to promoters, thereby regulating gene expression, development, stress responses, and pathogenicity in eukaryotes, including fungi[9, 10].

Traditional methods for studying histone modifications and chromatin-binding proteins have relied heavily on Chromatin Immunoprecipitation (ChIP) techniques, developed in the mid-1980s. The subsequent advent of ChIP, followed by high-throughput sequencing (ChIP-Seq), enables comprehensive identification of histone modifications or protein-binding sites across the whole genome. While ChIP-Seq has become a foundational approach for genome-wide chromatin analysis, it still has several limitations. The method requires large amounts of input material, may suffer from epitope masking, and often produces high background noise with low signal-to-noise ratios[11–14]. These challenges are particularly pronounced when studying filamentous fungi, where rigid cell walls and secondary metabolites reduce antibody accessibility and affect the efficiency of immunoprecipitation[15].

The recent development of Cleavage Under Targets and Tagmentation (CUT&Tag) offers a more efficient method for profiling chromatin features[16–18]. This technique employs a tethered Tn5 transposase, directed by specific antibodies, to simultaneously cleave DNA at target sites and integrate sequencing adapters. CUT&Tag-Seq offers several advantages over traditional ChIP-Seq, including higher-resolution mapping with lower cell input requirements (as few as 60 cells) and reduced background noise[19, 20]. While ChIP-seq remains the benchmark method for most chromatin studies, CUT&Tag offers unique advantages for studies that require live cell analysis or involve non-culturable samples, thereby providing a complementary approach to address the technical limitations of traditional methods. While CUT&Tag has been successfully applied to mammalian cells, plant cells, and yeast, its application in filamentous fungi has been limited due to technical challenges posed by the fungal cell wall.

In this study, we optimized a fungal CUT&Tag-Seq method (fCUT&Tag-Seq) for pathogenic/model fungi (e.g., *Verticillium dahliae*, *Neurospora crassa*), overcoming cell wall challenges via protoplast preparation and nuclear extraction. Formaldehyde crosslinking improved detection of low-abundance chromatin-binding proteins. Applied to histone modifications (H3K9me3, H3K4me3, etc.) and *SsGcn5*, our method yields high-quality data with low cell input, outperforming ChIP-Seq in signal-to-noise ratio and reproducibility in pathogenic fungi. fCUT&Tag-Seq streamlines fungal epigenetics research, reducing time and sample requirements while enhancing chromatin analysis.

## Results

### Optimization and Implementation of fCUT&Tag-Seq Method for Fungal Systems

We developed an optimized fCUT&Tag-Seq specifically adapted for fungal systems, building upon the traditional approach using the Vazyme Hyperactive Universal CUT&Tag Assay Kit for Illumina Pro (Cat. TD904). Our optimization addresses the unique challenges posed by fungal cell architecture while maintaining the high sensitivity and specificity characteristic of CUT&Tag methodology. The modified protocol incorporates several key steps for fungal applications **(Fig. 1).** To overcome the barrier presented by the fungal cell wall, we added a controlled enzymatic digestion step to generate protoplasts, followed by gentle nuclear extraction to preserve chromatin integrity. The isolated nuclei are then specifically captured by Concanavalin A-coated magnetic beads, which provide a stable platform for subsequent reactions while minimizing sample loss.

**Figure 1.**
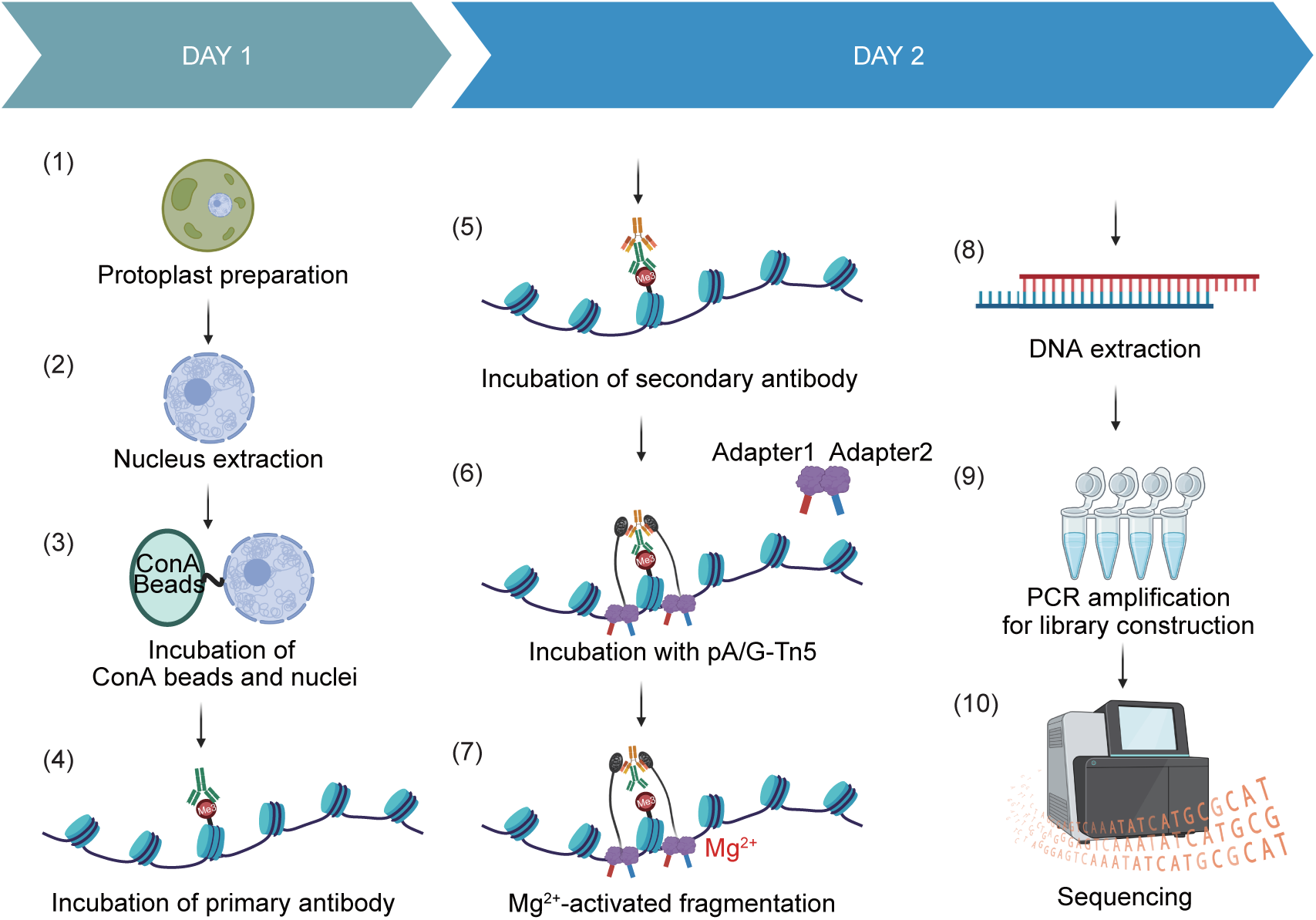
Schematic overview of the fCUT&Tag-Seq workflow for filamentous fungi. Illustration of the key steps in the optimized fCUT&Tag-Seq protocol in filamentous fungi, covering key steps from protoplast preparation to sequencing. The method involves digitonin permeabilization to enhance nuclear accessibility.

The workflow proceeds through several precisely controlled steps **(Fig. 1)**: (1) Nuclear membrane permeabilization is enhanced through carefully optimized digitonin treatment, facilitating antibody access while maintaining nuclear structural integrity. (2) Primary antibodies specific to the target histone modifications are introduced, followed by secondary antibodies that serve to amplify the detection signal. (3) The pA/G-Tn5 transferase complex is recruited to the antibody-bound locations through protein A/G interactions. (4) Upon Mg^2+^ activation, the tethered Tn5 simultaneously cleaves adjacent DNA and integrates sequencing adapters, enabling efficient library preparation through subsequent PCR amplification. After library construction, next-generation sequencing and data analysis can be performed to map the distribution pattern of histone modifications.

### fCUT&Tag-Seq Demonstrates Superior Quality Control Metrics and Target Enrichment Compared to ChIP-Seq in *V. dahliae*

To evaluate if we could use fCUT&Tag-Seq for profiling histone modifications in filamentous fungi, we first selected the plant pathogenic fungus *V. dahliae* as a model system. This pathogenic fungus, known for its broad host range, causes severe wilt, yellowing, and vascular browning in infected plants, often resulting in significant crop losses[21]. Given the known roles of H3K9me3 and H3K27me3 in gene silencing and pathogenicity[2, 22–24], we focused our analysis on these two modifications. To enhance our understanding of *V. dahliae*’s genetic background, we sequenced the whole genome of *V. dahliae* V592 strain using the Pacbio HiFi platform **(Fig. S1)**. Using the genome of this specific strain as the reference for mapping can help avoid errors introduced by inter-species genomic differences and also capture species-specific epigenetic modifications of genes, thereby providing more accurate information for the following analyses.

To evaluate the efficiency of our optimized fCUT&Tag-Seq method for profiling histone modifications in filamentous fungi, we conducted a comparative analysis in *V. dahliae* with other established methods. To validate the importance of protoplast extraction, we initially performed CUT&Tag experiments using spores from both wild-type and H3K9me3 methyltransferase mutant (VdΔ*kmt1*) strains to assess the genome-wide distribution of H3K9me3. However, this approach failed to generate a sufficient enrichment signal **(Fig. S2A)**. When comparing our approach with previously reported protocols specially developed for fungi (such as CUT&RUN for *Candida albicans* and CUT&Tag for *Saccharomyces pombe*), we attempted spheroplasting *V. dahliae* hyphae using 1.6 mg/mL Zymolyase[15, 25], incubating samples at 30°C for 30 min. However, this approach was unsuccessful as no spheroplasting occurred, with mycelial structures remaining intact **(Fig. S2B)**. Similarly, the application of the standard CUT&Tag protocol, designed for human cells, without prior protoplast extraction, yielded insufficient data[16]. Importantly, performing CUT&Tag directly on isolated protoplasts without nuclear extraction also failed to produce enrichment signals **(Fig. S2A)**, demonstrating that the nuclear extraction step is essential for successful CUT&Tag profiling in *V. dahliae*. Through further optimization, we determined that although using as low as 1 × 10^4^ cells was sufficient to show acceptable enrichment, 2.5 × 10LJ cells provided optimal data **(Fig. S2C).** Sequencing depths of approximately 1 Gb were sufficient to achieve the desired resolution **(Fig. S2D)**. These data collectively demonstrate the critical importance of protoplast preparation and nuclear extraction in our optimized fCUT&Tag-Seq protocol, enabling robust and high-quality profiling of histone modifications in filamentous fungi.

We then compared traditional ChIP-Seq data from the SRA database (Home - SRA - NCBI) and fCUT&Tag-Seq datasets based on multiple quality control metrics. Initial quality control analysis of fCUT&Tag-Seq libraries revealed characteristic nucleosomal ladder patterns through agarose gel electrophoresis, indicating successful fragmentation and library preparation **(Fig. 2A)**. Fragment length distribution analysis further confirmed the expected size distribution pattern typical of nucleosome-sized fragments **(Fig. 2B)**, validating the technical success of our protocol. Comparative analysis between fCUT&Tag-Seq and traditional ChIP-Seq demonstrated several key advantages of our optimized method. Most notably, fCUT&Tag-Seq exhibited a significantly higher percentage of reads mapped to peak regions and lower duplication rates (8.73%) compared to ChIP-Seq (88.45%), indicating substantially improved library complexity and data utilization efficiency **(Fig. 2C and 2D)**. Following alignment to the genome and subsequent filtering, we conducted a comprehensive analysis of quality metrics using ChIPQC. The fCUT&Tag-Seq data showed superior Signal Space Distance (SSD) scores, reflecting enhanced enrichment efficiency. Furthermore, the Fraction of Reads in Peaks (FRiP) consistently exceeded 7.8% in fCUT&Tag-Seq samples, with H3K9me3 modification showing particularly improved signal-to-noise ratios compared to ChIP-Seq **(Fig. 2D)**. And fCUT&Tag-Seq identified a greater signal intensity, indicating higher signal-to-noise **(Fig. 2E)**. Peak calling analysis revealed comparable patterns between fCUT&Tag-Seq and ChIP-Seq data, validating the biological relevance of our results. These comprehensive quality metrics demonstrate that our fCUT&Tag-Seq protocol not only meets standard quality benchmarks but also offers superior performance. The improved signal-to-noise ratio was particularly evident in genome-wide coverage plots, where fCUT&Tag-Seq showed more distinct enrichment patterns and clearer peak boundaries **(Fig. 2F)**. We have also performed strictly side-by-side comparisons of fCUT&Tag and ChIP-Seq in *V. dahliae*. The results showed that ChIP-Seq failed to show signal enrichment, while the fCUT&Tag assay yielded a strong H3K9me3 signal at the repeat region **(Fig. S2E)**. These comprehensive quality metrics demonstrate that our fCUT&Tag-Seq protocol not only meets standard quality benchmarks but also offers superior performance compared to traditional ChIP-Seq methods in terms of data quality, enrichment efficiency, and technical reproducibility in *V. dahliae*. The improved metrics suggest that this method provides a more reliable and efficient approach for studying histone modifications in *V. dahliae*.

**Figure 2.**
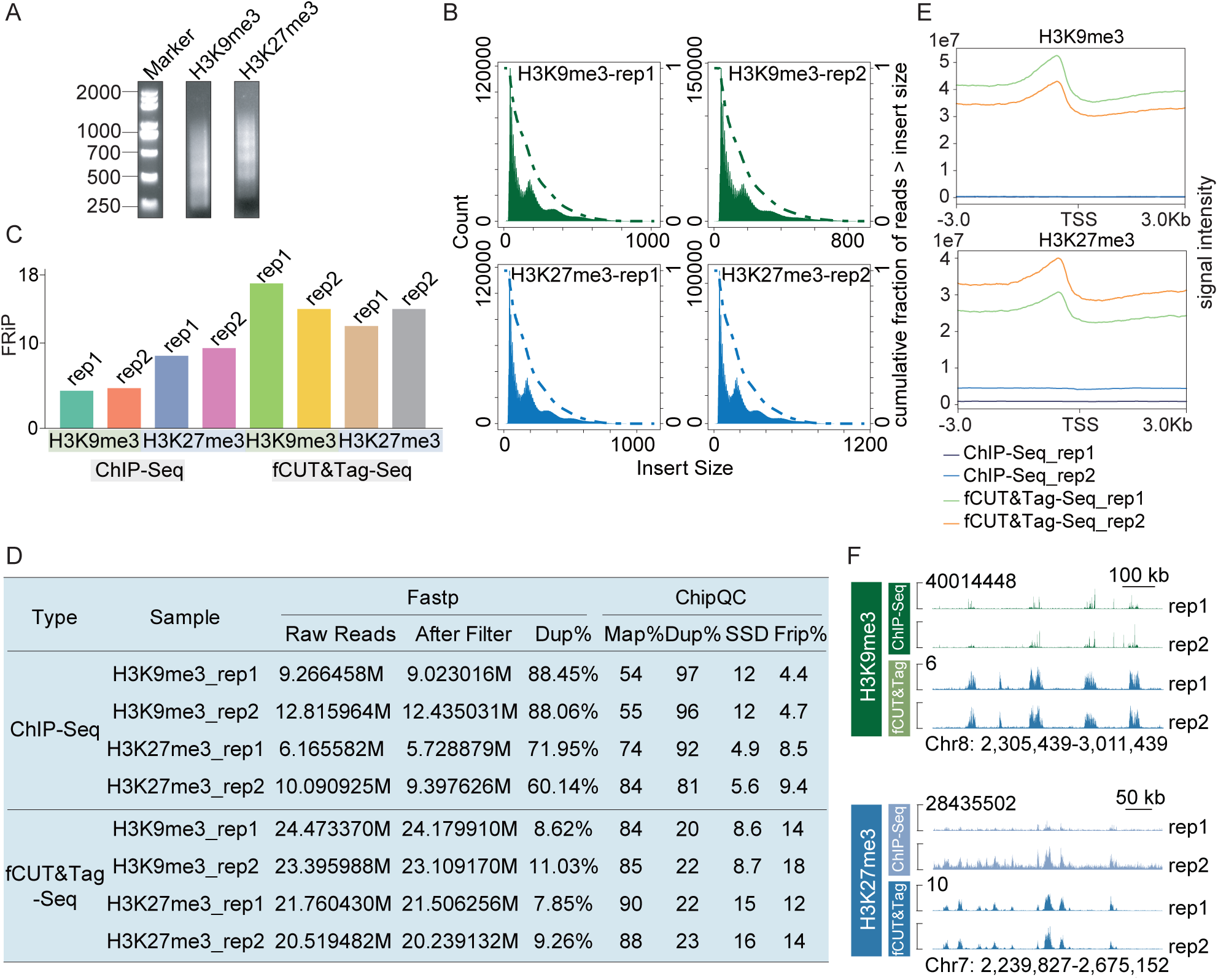
Comprehensive quality control analysis of fCUT&Tag-Seq compared to ChIP-Seq in *Verticillum dahliae*. **A.** Nucleosomal ladder pattern analysis of fCUT&Tag-Seq libraries by agarose gel electrophoresis. Libraries were generated from H3K9me3 and H3K27me3 histone modification profiling experiments in *Verticillum dahliae*. **B.** Fragment length distribution analysis of sequencing reads from fCUT&Tag-Seq and ChIP-Seq experiments. Green histograms represent H3K9me3 libraries (replicate 1 and replicate 2), while blue histograms show H3K27me3 libraries (replicate 1 and replicate 2). **C.** Bar plot showing the fraction of reads in peaks (FRiP) for the fCUT&Tag-Seq and ChIP-Seq datasets (replicates 1 and 2). **D.** Comparative analysis of key quality metrics between fCUT&Tag-Seq and ChIP-Seq methods. Quality assessment encompasses sequencing-level metrics (analyzed by Fastp), including total read counts, filtered read retention, and PCR duplicate rates, along with enrichment-specific parameters (evaluated by ChIPQC) measuring alignment efficiency (Map%), post-processing duplicates (Dup%), the uniformity of reads covering the entire genome (SSD), and Fraction of Reads in Peaks (Frip). **E.** Signal intensity profiles across peaks in fCUT&Tag-Seq versus ChIP-Seq datasets: the upper panel displays the H3K9me3 modification, while the lower panel shows the H3K27me3 modification. **F.** Genomic distribution analysis of identified peaks comparing fCUT&Tag-Seq and ChIP-Seq methods. The displayed region is randomly selected. The upper panel represents the H3K9me3 modification, and the lower panel shows the H3K27me3 modification.

### fCUT&Tag-Seq Method Efficiently Profiles Histone Modifications in *V. dahliae*

Following validation of our optimized protocol, we applied fCUT&Tag-Seq to investigate the genome-wide distribution of two key repressive histone modifications, H3K9me3 and H3K27me3, in *V. dahliae*. To evaluate the specificity of our method, we analyzed both wild-type strains and their corresponding methyltransferase mutants (VdΔ*kmt1* and VdΔ*ezh2)*. We incorporated spike-in controls for data normalization and used IgG antibodies as negative controls to account for background signal.

Genome-wide profiling revealed robust enrichment of both H3K9me3 and H3K27me3 modifications in wild-type strains **(Fig. 3A and 3B)**. H3K9me3 is a repressive mark mainly localized in constitutive heterochromatic regions, while H3K27me3 is a repressive mark responsible for facultative gene silencing, usually localized in euchromatin. In contrast, control experiments using non-specific IgG antibodies showed minimal signal across the genome, confirming the specificity of our approach **(Fig. S3A)**. Importantly, we observed significantly reduced levels of H3K9me3 and H3K27me3 in their respective methyltransferase mutants, VdΔ*kmt1* and VdΔ*ezh2*, consistent with the loss of these enzymatic activities **(Fig. 3A and 3B)**.

**Figure 3.**
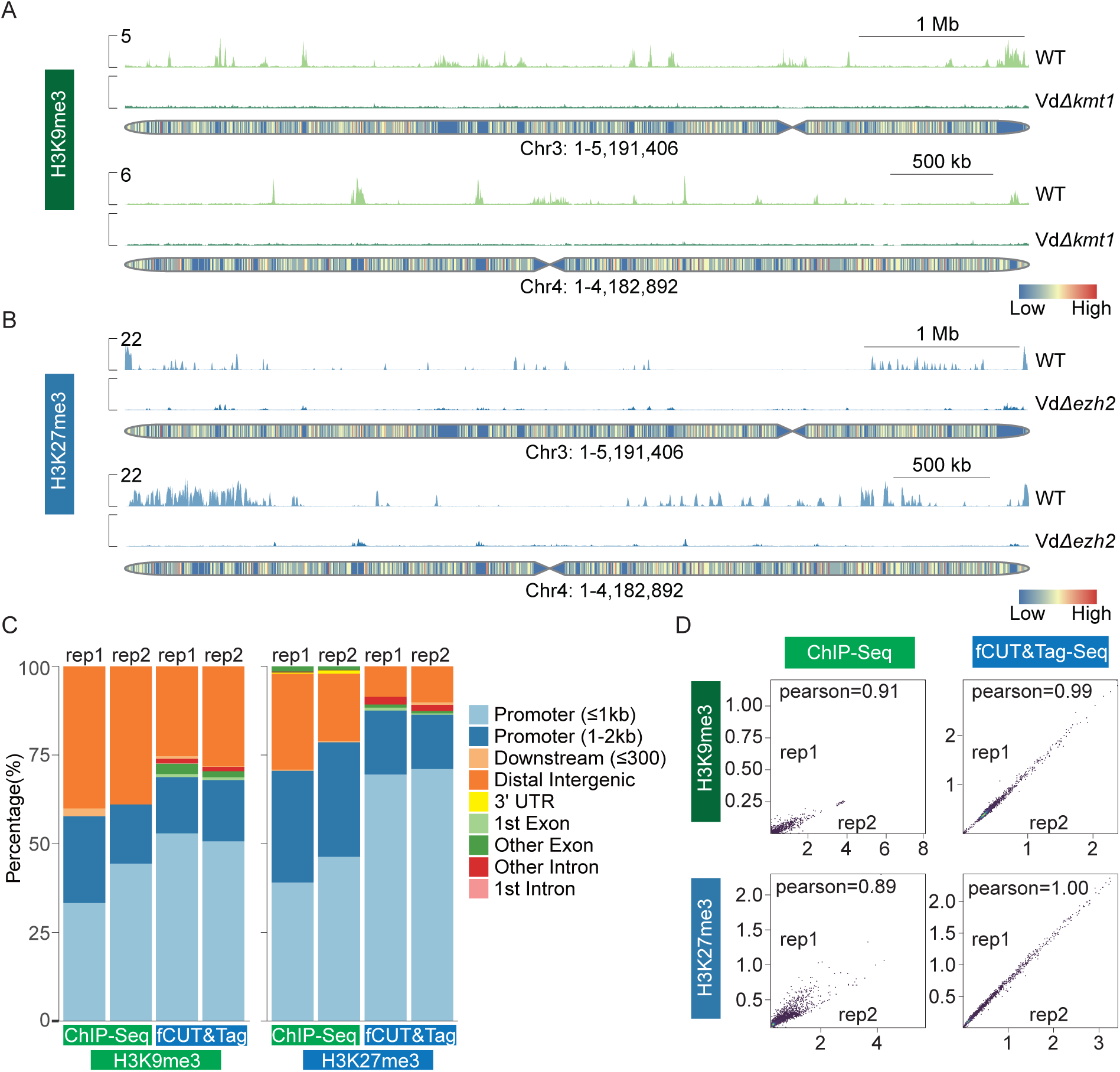
Genome-wide profiling of histone modifications in *V. dahliae* using fCUT&Tag-Seq. **A.** Genome browser view of H3K9me3 distribution in wild-type and VdΔ*kmt1* strains. **B.** Genome browser view of H3K27me3 distribution in wild-type and VdΔ*ezh2* strains. **C.** Comparative analysis of genomic feature distribution for H3K9me3 and H3K27me3 modifications between fCUT&Tag-Seq and ChIP-Seq approaches. The left panel shows the distribution of H3K9me3 peaks across different genomic elements, and the right panel shows H3K27me3 peak distribution. **D.** Pearson correlation analysis of signals between biological replicates for H3K9me3 and H3K27me3 modifications in ChIP-Seq (replicate 1 and replicate 2) and fCUT&Tag-Seq experiments (replicate 1 and replicate 2). Pearson correlations were calculated using deepTools, with the multiBamSummary tool followed by plotCorrelation. Read counts were divided into 1,000-bp bins across the genome for the analysis.

Analysis of the genomic distribution of these modifications revealed distinct patterns. Genomic feature distribution showed H3K9me3 peaks were predominantly located in gene promoter regions and distal intergenic spaces in constitutive heterochromatin, while H3K27me3 was mostly located in the gene promoter region in euchromatin **(Fig. 3C)**. These distribution patterns align with the known roles of these modifications in transcriptional regulation. A comparison of signal intensities between fCUT&Tag-Seq and ChIP-Seq data demonstrated superior enrichment in fCUT&Tag-Seq experiments, particularly across gene bodies and their 3-kb flanking regions **(Fig. S3B)**.

To assess the reproducibility of our modified method, we performed correlation analyses between biological replicates. fCUT&Tag-Seq demonstrated excellent reproducibility, with Pearson correlation coefficients of 0.99 and 1.0 for H3K9me3 and H3K27me3 modifications, respectively. These values exceeded those obtained from traditional ChIP-Seq experiments (0.91 and 0.89), highlighting the enhanced reliability of our optimized method **(Fig. 3D)**.

These results collectively demonstrate that our fCUT&Tag-Seq protocol enables highly specific and reproducible mapping of histone modifications in *V. dahliae*, providing a robust tool for investigating chromatin-based regulation in this important plant pathogen.

### fCUT&Tag-Seq Method Shows Broad Applicability Across Model Filamentous Fungi

To assess whether our fCUT&Tag-Seq protocol could be extended beyond *V. dahliae*, we tested it on two additional filamentous fungi: *N. crassa* and *F. graminearum*. Both species serve as valuable model organisms for studying fungal biology and pathogenicity, making them ideal candidates for validating broad applicability. Following the same experimental workflow, we focused on histone modifications H3K9me3 in *N. crassa*, and H3K4me3 in *F. graminearum*.

*N. crassa,* a member of the Ascomycota phylum, is widely recognized as a model organism due to its ease of cultivation and haploid genetics, which facilitate straightforward genetic analysis. Over the past few decades, it has been instrumental in studying various epigenetic modifications, including DNA methylation and gene silencing mechanisms[26–28]. We examined the levels of H3K9me3 in both the wild-type strain and the H3K9me3 methyltransferase mutant NcΔ*dim5*. Quality control assessments, including agarose gel electrophoresis and sequencing data analysis, confirmed that the fCUT&Tag-Seq libraries met the required standards **(Fig. 4A and 4B)**. The quality metrics indicated excellent performance for the *N. crassa* wild-type samples **(Fig. 4C)**. Notably, the peak distribution patterns observed were consistent with those identified through ChIP-Seq, yet fCUT&Tag-Seq demonstrated higher signal intensity **(Fig. 4D)**. The correlation between fCUT&Tag-Seq and ChIP-Seq data reached 0.73 **(Fig. 4E)**, further validating the reliability of our method. Additionally, biological replicates exhibited strong correlation, indicating high reproducibility **(Fig. 4F)**. Comparative analysis revealed that H3K9me3 was significantly enriched in the wild-type strain, while the NcΔ*dim5* mutant displayed markedly lower modification levels **(Fig. 4G)**. As expected, H3K9me3 modifications were predominantly located in distal intergenic regions and promoter regions **(Fig. 4H)**.

**Figure 4.**
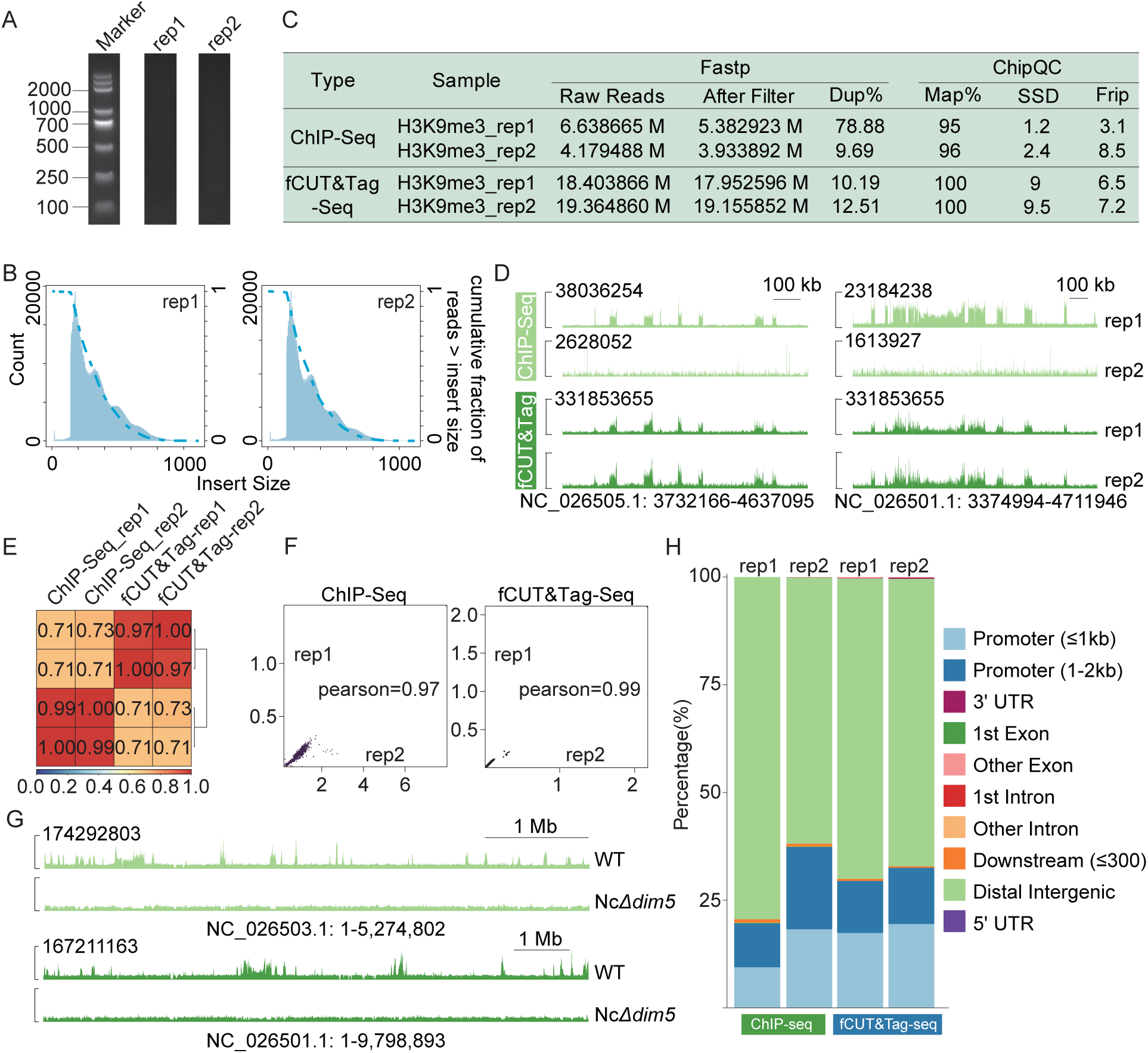
Application of fCUT&Tag-Seq for histone modifications analysis in *N. crassa*. **A.** Nucleosomal ladder pattern analysis of fCUT&Tag-Seq libraries by agarose gel electrophoresis. Libraries were generated from two replicates of H3K9me3 profiling experiments in *N. crassa*. **B.** Fragment length distribution analysis of sequencing reads from fCUT&Tag-Seq experiments. Histograms display the expected nucleosomal fragmentation patterns for two replicates. **C.** Comparative analysis between fCUT&Tag-Seq and ChIP-Seq data for H3K9me3 modification in *N. crassa*. **D.** Genome browser view of peak distribution analysis comparing fCUT&Tag-Seq and ChIP-Seq results. The upper panel shows H3K9me3 enrichment in ChIP-Seq, and the lower panel displays the modification distribution from fCUT&Tag-Seq. **E.** Correlation matrix analysis between fCUT&Tag-Seq and ChIP-Seq data, indicating strong agreement. Pearson correlations were calculated using deepTools (multiBamSummary followed by plotCorrelation), based on read counts divided into 1000-bp bins across the genome. **F.** Pearson correlation analysis of H3K9me3 signals between biological replicates in ChIP-Seq and fCUT&Tag-Seq. **G.** Genome browser view of the H3K9me3 peaks accross representative genomic regions in wild-type and NcΔ*dim5* strains in *N. crassa*. **H.** Feature distribution comparison of H3K9me3 modification between ChIP-Seq and fCUT&Tag-Seq.

Similar results were obtained for *F. graminearum*, the causative agent of fusarium head blight in wheat[29]. In addition to analyzing H3K9me3 and H3K27me3 modifications, we also assessed H3K4me3 levels, which are associated with gene activation. The sequencing quality metrics of fCUT&Tag-Seq for *F. graminearum* were superior to those of ChIP-Seq, with a greater percentage of reads mapping to peak regions **(Fig. 5A and 5B)**. The Signal Space Distance (SSD) score and the percentage of reads in peaks (Rip%) reached 21 and 46, respectively **(Fig. 5B)**. These findings confirm the robustness of our method, as the wild-type strain exhibited significant enrichment **(Fig. 5C)**. Notably, peak enrichment was predominantly observed in the promoter regions (96.51%) **(Fig. 5D)**. The Pearson correlation coefficient was 0.99, indicating a strong correlation between the sample groups **(Fig. 5E)**.

**Figure 5.**
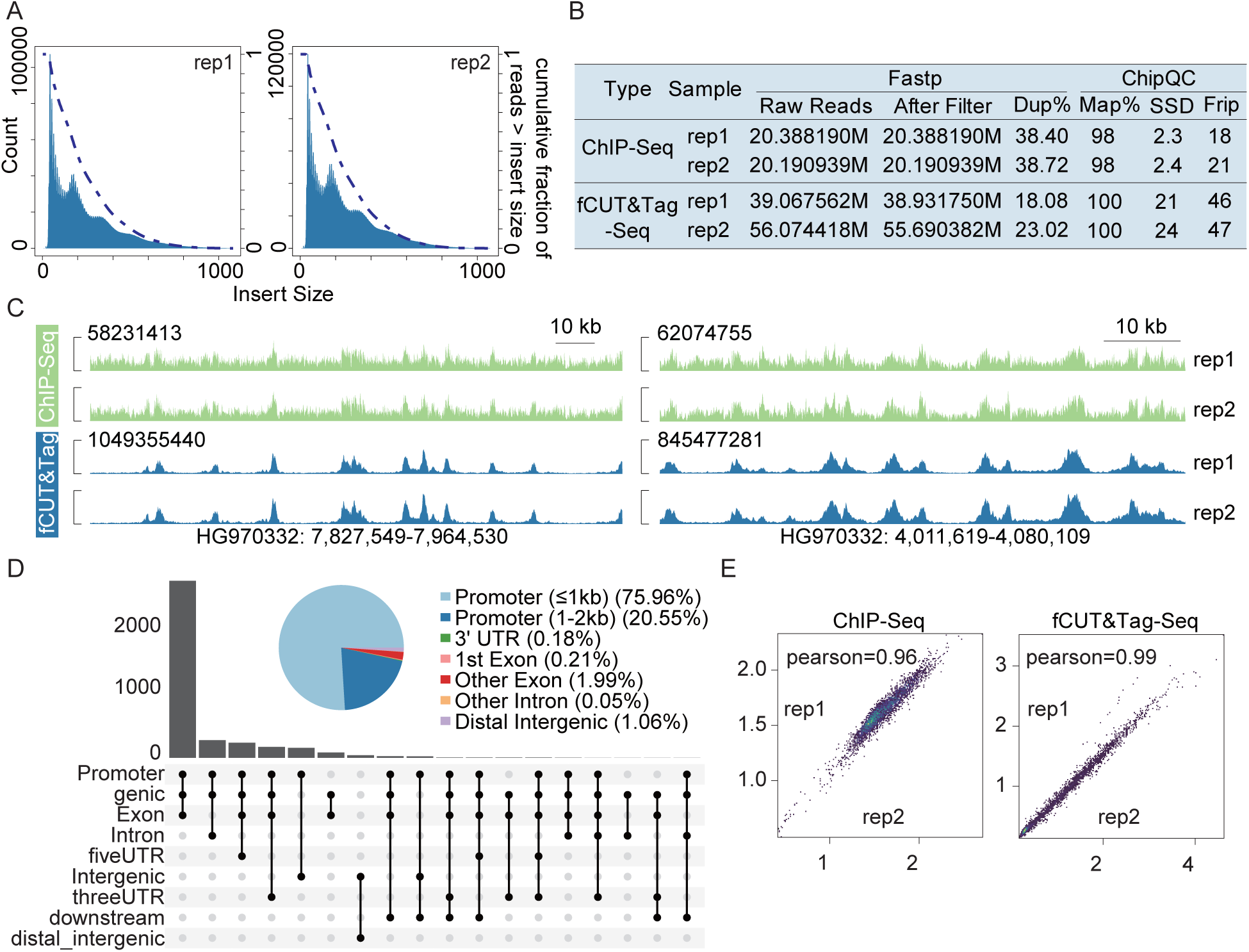
Genome-wide profiling of H3K4me3 modification in *F. graminearum* using fCUT&Tag-Seq analysis. **A.** Fragment length distribution of sequencing libraries for fCUT&Tag-Seq. The left panel shows the fragment length distribution for the H3K4me3 library in replicate 1, and the right panel shows the distribution for replicate 2. **B.** Comparative analysis of key quality metrics for ChIP-Seq and fCUT&Tag-Seq datasets. **C.** Genome browser view of H3K4me3 modification peak distribution analysis comparing fCUT&Tag-Seq and ChIP-Seq results. The genome regions were randomly selected. Green peaks represent ChIP-Seq results, and blue peaks represent fCUT&Tag-Seq results. **D.** The pie chart and UpSet plot illustrate the distribution of peaks across different genomic features. The results showing that H3K4me3 is predominantly enriched in promoter regions. Distinct colors represent different genomic features. **E.** Pearson correlation coefficients between sample groups for ChIP-Seq and fCUT&Tag-Seq. The scatter plot displays normalized read counts in 1,000-bp bins across the genome.

Overall, these results demonstrate that our fCUT&Tag-Seq method is not only effective for detecting histone modifications in *V. dahliae*, but also applicable to other model fungi, including *N. crassa* and *F. graminearum*. This versatility underscores the potential of fCUT&Tag-Seq as a powerful tool for studying epigenetic modifications across a broad range of fungal species.

### fCUT&Tag-Seq Method Enables Profiling of Chromatin-Binding Proteins

To extend the utility of our optimized fCUT&Tag-Seq protocol beyond histone modifications, we adapted the method to profile chromatin-binding proteins, which play crucial roles in transcriptional regulation. Given the typically lower abundance of chromatin-binding proteins compared to histones, we incorporated an additional formaldehyde crosslinking step before protoplast preparation. This modification strengthens protein-chromatin interactions, thereby improving detection sensitivity. Following fragmentation, we performed sequential decrosslinking, protease digestion, and RNase treatment to ensure efficient DNA purification and library preparation **(Fig. 6)**.

**Figure 6.**
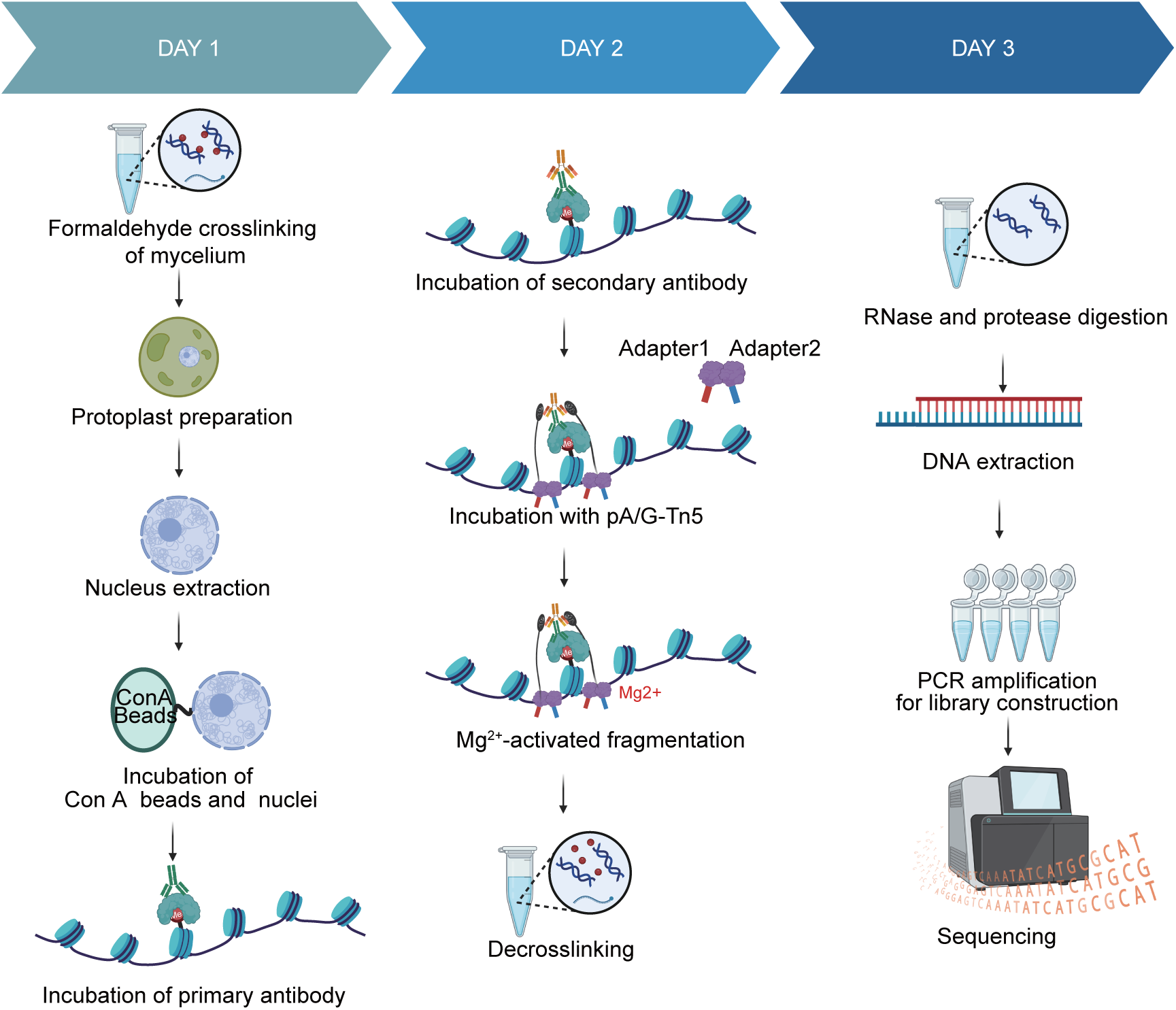
Schematic overview of the optimized fCUT&Tag-Seq method for detecting chromatin-binding proteins. The workflow diagram illustrates the key steps of the fCUT&Tag-Seq protocol for analyzing chromatin-binding proteins in fungi, including protoplast preparation, formaldehyde crosslinking, nuclear extraction, and subsequent fCUT&Tag-Seq procedures.

To validate this adapted fCUT&Tag-Seq method, we investigated the histone acetyltransferase SsGcn5 in *S. scitamineum*, a dimorphic pathogenic fungus responsible for sugarcane smut. Sugarcane smut is one of the most significant diseases affecting global sugarcane production worldwide. The disease poses substantial economic threats, highlighting the urgent need for effective prevention and control strategies against *S. scitamineum*[30–32]. It has been reported that histone acetylation may play a critical role in fungal pathogenesis[33–36]. To explore the relationship between histone acetylation and fungal pathogenicity, we screened possible histone acetyltransferase candidate genes and introduced a Flag epitope tag into the histone acetyltransferase Gcn5 in two *S. scitamineum* strains with different mating types (JG35 and JG36), using our previously reported method[37].

Quality control analysis of the resulting fCUT&Tag-Seq libraries revealed proper DNA fragmentation patterns by agarose gel electrophoresis and expected nucleosomal size distributions in sequencing reads **(Fig. 7A and 7B)**, confirming successful library preparation. Comprehensive quality metrics and genome-wide profiling demonstrated robust enrichment of SsGcn5-3Flag binding sites across the *S. scitamineum* genome **(Fig. 7C and 7D)**. Notably, comparative analysis revealed a strong correlation between SsGcn5 binding sites and regions of H3K18 acetylation **(Fig. 7E)**, consistent with the known catalytic activity of Gcn5. Feature distribution analysis showed that SsGcn5 binding sites were predominantly located in promoter regions (85.93% of total peaks) **(Fig. 7F)**, suggesting a primary role in transcriptional regulation. The high reproducibility of our method was demonstrated by the near-perfect Pearson correlation coefficient (approximately 1.0) between biological replicates **(Fig. 7G)**.

**Figure 7.**
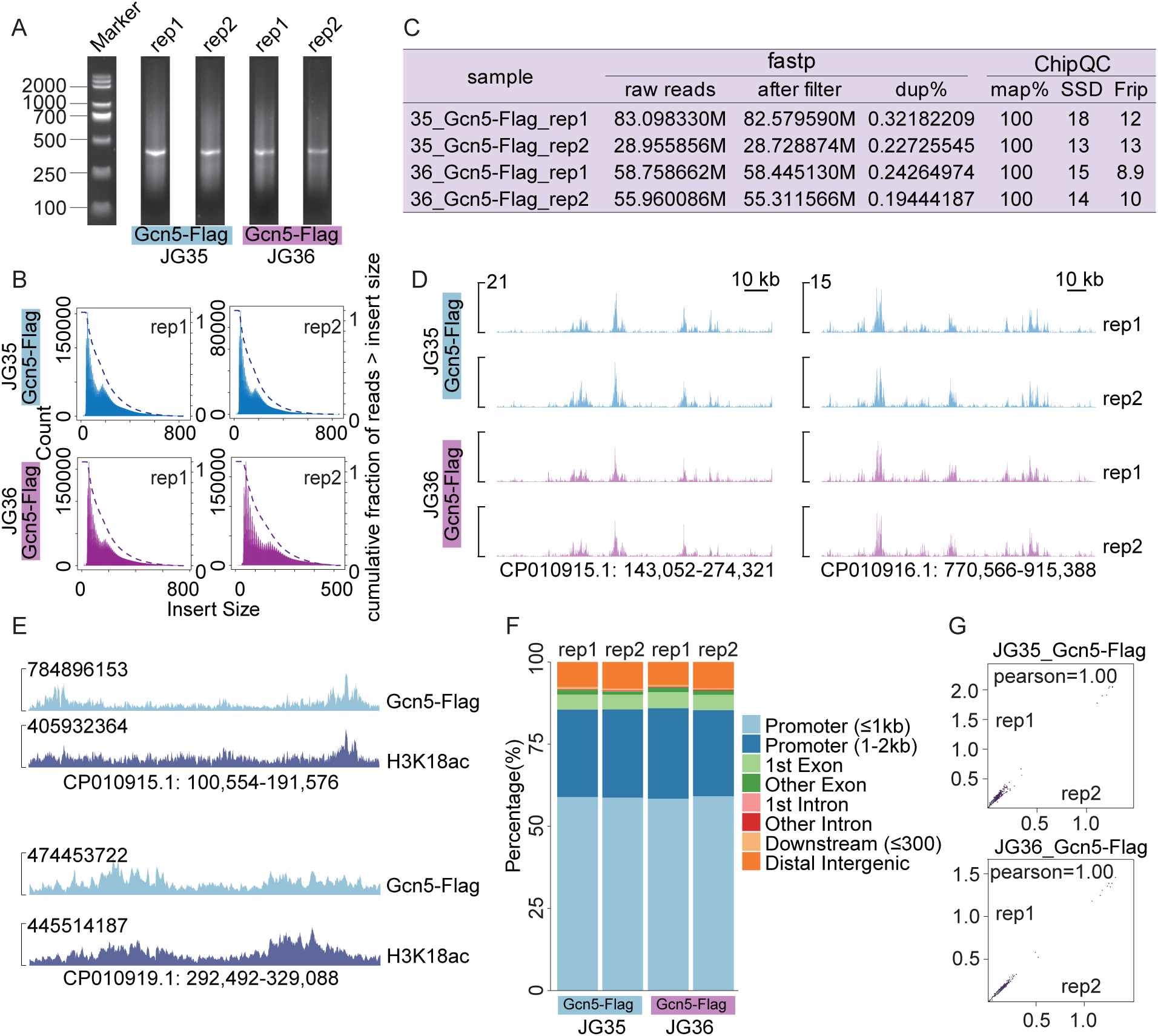
Genome-wide profiling of SsGcn5 and its correlation with H3K18ac modification in *S. scitamineum* using fCUT&Tag-Seq. **A.** Quality assessment of fCUT&Tag-Seq libraries by agarose gel electrophoresis. DNA fragment distributions are shown for two biological replicates of JG35 (lanes 1-2) and JG36 (lanes 3-4) mating types. **B.** Fragment length distribution analysis of sequencing reads demonstrating expected nucleosomal patterns. Blue and pink histograms represent Gcn5-Flag distributions in JG35 and JG36 strains, respectively. **C.** Comprehensive quality metrics for fCUT&Tag-Seq datasets in DNA-binding protein enrichment in JG35 and JG36 strains. **D.** Genome browser view of SsGcn5-Flag binding sites identified by fCUT&Tag-Seq. Genome regions were selected randomly. **E.** Correlation analysis between SsGcn5-Flag binding sites and H3K18ac modification regions, showing the functional relationship between histone acetyltransferase localization and its catalytic activity. **F.** Genomic feature distribution analysis of SsGcn5-Flag peaks in JG35 and JG36 strains, showing the relative enrichment across different genomic elements (promoters, gene bodies, intergenic regions). **G.** Correlation analysis between biological replicates for SsGcn5-Flag fCUT&Tag-Seq experiments.

Our results confirmed that the fCUT&Tag-Seq method not only excels in detecting histone modifications, but also effectively profiles chromatin-binding proteins in fungi. By enhancing the detection of low-abundance proteins, this method also provides a powerful tool for exploring chromatin-binding proteins in regulating gene expression and chromatin dynamics across a broad range of fungal species.

## Discussion

Over prolonged evolutionary periods, plant pathogenic fungi have developed intricate strategies to penetrate and colonize their hosts. Precise control of pathogenic gene expression is pivotal to these infection processes, influencing fungal development, environmental stress responses, and virulence[38]. Emerging evidence highlights the significance of epigenetic regulation, including histone methylation, histone acetylation, and chromatin remodeling, in orchestrating transcriptional reprogramming in plant pathogens[39–41]. However, until recently, the available tools for epigenetic research in diverse fungal systems were still very limited, making advanced investigations into fungal chromatin biology challenging.

In this study, we optimized and extended CUT&Tag-Seq to several important fungal species, including *V. dahliae*, *F. graminearum*, *N. crassa*, and *S. scitamineum*. Our fCUT&Tag-Seq method effectively addresses the unique obstacles posed by fungal cell walls by integrating protoplast preparation and gentle nuclear extraction. Additionally, a formaldehyde crosslinking step was introduced to strengthen DNA-protein interactions, particularly beneficial for profiling low-abundance chromatin-binding proteins.

Our data revealed that the fCUT&Tag-Seq method accurately and sensitively detects histone modifications (H3K9me3, H3K27me3, H3K4me3, and H3K18ac) as well as chromatin-binding proteins in these fungi. We also performed genome-wide profiling of H3K9me3 and H3K27me3 distributions, revealing distinct compartmentalization patterns characteristic of fungal epigenomes. Our analyses demonstrate that H3K9me3 predominantly marks constitutive heterochromatin regions, showing strong enrichment at pericentromeric and telomeric domains, consistent with its role in silencing repetitive elements[42–44]. In contrast, H3K27me3 preferentially marks facultative heterochromatin, regulating transcriptionally dynamic regions that modulate stress responses and developmental transitions[45, 46]. Its reversible deposition enables rapid gene expression remodeling in response to environmental cues, such as host infection or nutrient availability[47].

As epigenetic modifications play crucial functional roles during fungal infection processes. Taking advantage of the compositional differences between plant and fungal cell walls, we would be able to isolate fungal protoplasts in planta through enzymatic digestion directly in the near future. Using the fCUT&Tag-Seq method, we may profile fungal-specific epigenetic modifications during host infection, thereby studying their critical associations with fungal pathogenicity. Notably, the minimal cellular input requirement of this methodology (potentially as few as 10,000 cells) may offer future applications for studying unculturable fungal species.

While recent studies have established robust ChIP-Seq protocols for model fungi like *N. crassa* (e.g., datasets SRR1566112, SRR12229310) [48, 49], our comparative analyses revealed measurable differences in H3K9me3/H3K27me3 detection sensitivity and reproducibility between established methods and our fCUT&Tag-Seq approach. More importantly, our method mainly lies in its consistency across non-model fungi, where traditional ChIP-Seq is particularly challenging.

Although our fCUT&Tag-Seq method has demonstrated significant improvements in data quality and sensitivity, it is still important to acknowledge the limitations. Challenges remain in enriching low-abundance proteins and chromatin-binding proteins located in heterochromatin regions, where accessibility to antibodies is largely reduced. Future studies should focus on further optimizing methods for relaxing heterochromatin to enhance the detection of these proteins.

## Materials and Methods

### Fungal Strains and Culture Conditions

The following fungal strains were used in this study: *V. dahliae* strain V592 (isolated from cotton in Xinjiang, China)[50, 51], *N. crassa* 87-3 and NcΔ*dim5* (provided by Xiao Liu) [52], *S. scitamineum* strains JG35 and JG36 (provided by Shan Lu) [53], and *F. graminearum* wild-type strain (provided by Guangfei Tang and Haoxue Xia)[54]. All strains were maintained and cultured according to standard protocols specific to each species.

### Generation of Mutant and Tagged Fungal Strains

For *V. dahliae*, *VdKMT1* (*VDAG_07826*) and *VdEZH2* (*VDAG_00983*) deletion strains (VdΔ*kmt1* and VdΔ*ezh2*) were constructed as previously reported[55]. 1 kb genomic sequences upstream and downstream of target genes were amplified from V592 DNA with the following primer pairs: VdKMT1-A/ VdKMT1-a, VdKMT1-d/ VdKMT1-B, VdEZH2-A/ VdEZH2-a, VdEZH2-d/ VdEZH2-B (listed in Table S1) using the Phanta Max Super-Fidelity DNA Polymerase (P505, Vazyme) and cloned into the pGKO-HPT/pGKO-NAT vector *via* homologous recombination. Sequence-verified vectors were then used to transform *V. dahliae* by *Agrobacterium tumefaciens-*mediated transformation (ATMT) system. *V. dahliae* transformants were selected on potato dextrose agar (PDA) plates supplemented with 5-fluoro-2’deoxyuridine (5FU) and hygromycin B / Nourseothricin. Putative transformants were screened using PCR to confirm the successful deletion of the target genes with primer pairs V1/Hpt-R and V2/Hpt-F. The primers used for construction are listed in Table S1.

For *S. scitamineum*, the SsGcn5-Flag strain was constructed using homologous recombination. The vector pEX1-GAP-Flag-HPT, carrying a Flag tag, was modified from pEX1-GAP-HPT. Approximately 1.5 kb of upstream (5’) and downstream (3’) flanking sequences near the target protein’s stop codon (excluding the stop codon) were amplified from the wildtype strains JG35 and JG36 using primer pairs: SsGcn5-tg-A/SsGcn5-a and SsGcn5-d/SsGcn5-B, respectively (Table S1). Two truncated fragments of the hygromycin resistance gene HPT (HPT-up and HPT-down) were amplified from the pEX1-GAP-Flag-HPT plasmid with the primer pairs HPT-LB-F/HPT-LB-R and HPT-RB-F/HPT-RB-R. Overlap PCR was employed to fuse the upstream sequence with HPT-up (primers SsGcn5-tg-A/HPT-LB-R) and the downstream sequence with HPT-down (primers HPT-RB-F/SsGcn5-B). The products were introduced into *S. scitamineum* strains JG35 and JG36 via PEG-mediated protoplast transformation. Transformants were selected on YEPSA medium containing 150 μg/mL hygromycin B and verified by PCR using primer pairs SsGcn5-C(V1)/HPT-LB-R and SsGcn5-D(V2)/HPT-RB-F.

### fCUT&Tag-Seq for Histone Modification

To profile histone modifications using fCUT&Tag-Seq, protoplasts were prepared by digesting mycelium (*V. dahliae, N. crassa,* and *F. graminearum*) or spores (*S. scitamineum*) with 10 mL of enzyme solution (0.25 g VinoTaste^®^Pro enzyme in 12.5 mL 0.7 M NaCl for *V. dahliae*, *N. crassa,* and *S. scitamineum*; 0.1 g Diselease, 0.2 g Celulase, and 0.2 g Lysozyme were diluted with 0.7 M NaCl to 20 mL for *F. graminearum*). *F. graminearum* was incubated at 30°C with shaking at 85 rpm, while the others were incubated at 33°C at 60 rpm. Digestion times were 3 h for *V. dahliae* mycelium, *F. graminearum* mycelium, and *N. crassa* mycelium, and 30 min for *S. scitamineum* spores. Protoplasts were filtered through Miracloth (475855-1R; Merck), and centrifuged for 10 min at 3800 rpm (*V. dahliae* and *N. crassa*), 2200 rpm (S*. scitamineum*), or 5000 rpm (*F. graminearum*) at 25LJ. Protoplasts were resuspended in 10 mL of 0.7 M NaCl and then counted using a hemocytometer.

Nuclei were extracted from 1 million protoplasts per 1.5 mL tube by centrifugation at 3000 rpm for 5 min at 25 °C, followed by resuspension in 100 μL nuclear extract buffer (TD904; Vazyme), incubation on ice for 10 min, and a second centrifugation at 2500 rpm for 6 min at 25 °C. The supernatant was discarded, and nuclei were resuspended in 100 µL of wash buffer (TD904; Vazyme).

Nuclei were adsorbed by mixing activated Concanavalin A-coated magnetic beads (TD904; Vazyme) with 250,000 resuspended nuclei and 75 µl of wash buffer (TD904; Vazyme) in 200μL 8-row tubes, followed by incubation at room temperature for 10 min with 2-3 inversions.

For antibody binding, tubes were placed on a magnetic rack, the supernatant was discarded, and 50 μL of antibody buffer (TD904; Vazyme) was added to resuspend the bead-nucleus complex. Primary antibodies (1 μL each) against H3K9me3 (Cat39161; Active Motif), H3K27me3(Cat39155; Active Motif), or H3K4me3 (Cat39060; Active Motif) were added, according to the ratio used. After mixing by inverting the tube, centrifuge immediately, then place in a 4℃ refrigerator for overnight incubation under static conditions (no rotation).

The next day, the beads were centrifuged briefly, placed on a magnetic rack, and the supernatant was discarded. 0.5 µL of secondary rabbit antibody (Ab207; Vazyme) in 49.5 µL of Dig wash buffer (792 μL Wash Buffer, 8 μL 5% Digitonin) was added and incubated with rotation (9-11 rpm/min) at room temperature for 1h. Complexes were washed three times with 200 µL of Dig wash buffer.

For transposase incubation, 2 µL of Hyperactive pA/G-Transposon (TD904; Vazyme) in 98 µL Dig-300 buffer (100 μL 10 × Dig-300 Buffer, 2 μL 5% Digitonin, 20 μL 50 × protease inhibitor, 878 μL ddH_2_O) was added to each reaction and incubated with gentle rotation (9-11 rpm/min) at room temperature for 1 h. Subsequently, after instantaneous centrifugation and placing on a magnetic rack, supernatant was discarded and washed three times using 200 µL Dig-300 buffer.

For fragmentation, 10 µL of TruePrep Tagment Buffer L (TD904; Vazyme) in 40 µL Dig-300 buffer was added to each reaction and incubated at 37°C for 1 h.

DNA was extracted by adding 2 µL of 10% SDS (TD904; Vazyme) and 0.5 µL of DNA spike-in (TD904; Vazyme), incubating at 55°C for 10 min, transferring supernatant to a new tube, adding 50 μL activated DNA extract beads (TD904; Vazyme), incubating for 20 min at room temperature with mixing, washing twice with 200 μL 1× B&W buffer (TD904; Vazyme), drying, and resuspending in 30 µL of RNAase-free ddH_2_O.

Libraries were constructed by PCR (50 μL reaction: 15μL bead-bound DNA, 25 μL 2× CUT&Tag Amplification Mix [TD904; Vazyme], 5μL N5XX Primer [TD202; Vazyme], 5μL N7XX Primer [TD202; Vazyme]) following Illumina’s adaptor pooling guide. PCR conditions: 72°C for 3 min, 95°C for 3 min; then 9-20 cycles of 98°C for 10 s, 60°C for 5 s, 72°C for 1 min; hold at 4°C.

PCR products were purified by adding 100 μL VAHTS DNA clean beads (N411; Vazyme), incubating for 5 min, washing twice with 200 μL 80% ethanol, drying, resuspending in 22 μL RNase-free ddHLJO, and incubating for 5 min before quality control and sequencing.

### fCUT&Tag-Seq for chromatin-binding proteins

To profile lower-abundance chromatin-binding proteins, a formaldehyde crosslinking step was added. Approximately 1 g of fungal spores was fixed with 9 mL of 0.1% formaldehyde for 8 min at room temperature, quenched with 1 mL of 2 M glycine for 5 min at room temperature. The cross-linked cells were washed three times with 20 mL of 0.7 M NaCl.

Protoplast formation, nuclei capture with ConA-beads, antibody incubation, and pA/G-Tn5 transposome assembly followed the histone modification protocol. After fragmentation, crosslinks were reversed by incubating overnight at 65°C with 3 μL 5 M NaCl. RNA was removed by RNase A treatment (37°C for 30 min), proteins by Proteinase K (53°C for 1 h), and proteases by inhibitors (room temperature for 5 min). Libraries were amplified by PCR as described.

### ChIP-Seq

ChIP assays were performed as previously described[56–58]. Mycelia were cross-linked with 1% formaldehyde solution (0.4 M sucrose, 10 mM Tris-HCl [pH 8.0], 1 mM EDTA, 1 mM PMSF, 1% formaldehyde) at room temperature for 15 minutes, with the flask inverted every 2–3 minutes. To stop cross-linking, glycine was added to a final concentration of 0.2 M, followed by incubation for 5 minutes. Fixed cells were then washed with 1× PBS, cryomilled in liquid nitrogen, and lysed in lysis buffer (50 mM HEPES [pH 7.5], 137 mM NaCl, 1 mM EDTA, 0.1% SDS, 1% Triton X-100, 0.1% sodium deoxycholate, 1 mM PMSF, 1 μg/mL leupeptin, 1 μg/mL pepstatin A). Chromatin was sonicated to an average DNA fragment size of ∼500–1000 bp. After sonication, samples were centrifuged at 10,000 ×g and 4 °C for 15 minutes to collect the supernatant. For immunoprecipitation, 2 μL of anti-H3K9ac antibody (Cat. No. 39161; Active Motif) was incubated with aliquots of clarified cell lysates (containing 2 mg protein each) at 4 °C overnight with rotation. After overnight incubation, 50 μL of SS DNA/Protein G Sepharose (pre-treated three times with lysis buffer) was added, followed by further incubation at 4 °C for 2 hours. Sepharose beads were collected and washed once with 1 mL of each of the following buffers: high-salt wash buffer (lysis buffer supplemented with 0.5 M NaCl), 1× LNDET (0.25 M LiCl, 1% NP40, 1% deoxycholate, 1 mM EDTA, 10 mM Tris [pH 8]), and 50 mM Tris-HCl [pH 8], 1 mM EDTA. Immune complexes were eluted from the beads using freshly prepared elution buffer (1% SDS, 0.1 M NaHCOLJ) via two rounds of elution at room temperature (250 μL per elution).

### fCUT&Tag-Seq Data Analysis

Raw ChIP-Seq data were downloaded from the SRA database using the SRA Toolkit (version 3.1.1). Raw data of ChIP-Seq and fCUT&Tag-Seq were first processed using fastp (0.23.4)[59] to remove adapters and low-quality sequences. The cleaned reads were then aligned to the spike-in sequences using BWA (0.7.18)[60]. After alignment, SAMtools (version 1.21)[61] was used to count the number of successfully mapped reads. The sample with the fewest mapped reads was selected as the baseline for spike-in normalization. The normalized data were then re-aligned to the reference genome using BWA (0.7.18), with reads having a quality score below 20 being filtered out using samtools (1.21). The alignment results were then processed using Picard (<u>Picard Tools, developed by the Broad Institute</u>) for sorting and deduplication. Peak calling was performed using MACS2 (2.2.9.1)[62], with IgG controls used to correct for background. The “-B” flag was employed to generate histone modifications and control lambda .bdg files, which were subsequently compared using the bdgcmp function in MACS2. ChIPQC (1.42.0, <u>Bioconductor -</u> <u>ChIPQC</u>) was applied for quality control of the fCUT&Tag analysis. Utilize Spark.py (2.6.2)[63] to generate a genomic browser view, incorporating the -gs parameter to ensure all groups are autoscaled to a unified y-axis. Peak annotation was performed using ChIPseeker (1.42.0)[64] to assign genomic features to the identified peaks. Correlation analysis was carried out using the multiBigwigSummary and plotCorrelation functions of deepTools (3.3.5)[65], while heatmaps were generated using the computeMatrix and plotHeatmap functions of deepTools (3.3.5).

## Supporting information

**Fig. S1 *V. dahliae* V592 genome assembly presentation.** (A) The output quality of sequencing data. Sample: sample name; SeqNum: number of sequences; SumBase: total number of databases; N50Len: N50 length of sequencing data; N90Len: length of sequencing data N90; MeanLen: mean length of sequencing sequence; MaxLen: maximum length of sequencing sequence. (B) Assembly result statistic. Length: length of the sequence after concatenation; GC (%): GC content of the concatenated sequence. (C) Coding gene prediction results. (D) The CIRCOS plot of the genome.

**Fig.S2 Optimization and comparative evaluation of the fCUT&Tag-Seq protocol for *V. dahliae*.** (A). Genome browser view of H3K9me3 distribution in wild-type and VdΔ*kmt1* strains using nucleus, spores or protoplast. (B) Comparison of the state of protoplasts extracted by different methods. The upper panel showed that no spheroplasting state was observed, extracted by zymolyase described in the CUT&RUN approach for *Candida albicans* and the CUT&Tag method for *Saccharomyces pombe*, and the lower panel showed the protoplast state using the optimized fCUT&Tag-Seq technique. The images were taken by a confocal microscope (Leica TCS SP8). Scale bar: 25 µm. (C) Genome browser view of H3K9me3 distribution in wild-type and VdΔ*kmt1* strains using different cell numbers. (D) Genome browser view of H3K9me3 distribution in wild-type and VdΔ*kmt1* strains using different sequencing volumes. (E) Genome browser view of H3K9me3 distribution in wild-type and VdΔ*kmt1* strains using fCUT&Tag-Seq and ChIP-Seq, respectively. The Y-axis values in ChIP-seq tracks represent normalized read coverage depth (in reads per million, RPM), calculated by scaling raw read counts to 10LJ.

**Fig.S3 Genome-wide profiling of nonspecific IgG signals and histone modification patterns in *V. dahliae* using fCUT&Tag-Seq.** (A) Genome browser view of the H3K9me3, H3K27me3, and nonspecific IgG antibody signals across representative genomic regions in the wild-type strain. Data were presented at the same scale as specific antibody signals. (B) Heatmap analysis of H3K9me3 and H3K27me3 signals near protein-coding genes in fCUT&Tag-Seq and ChIP-Seq. Scale regions were 3,000 bp upstream of the translation starting site (TSS), 3,000 bp downstream of the translation end site (TES), and a 1,000-bp region on the gene body. The lengths were plotted using the computeMatrix and plotHeatmap tools in deepTools.

**Fig.S4 Original images of Figure 2A**, **Figure 4A and Figure 7A**. (A) Original images of Figure 2A. (B) Original images of Figure 4A. (C) Original images of Figure 7A.

**Table.S1 Primers used in this study**.

## Acknowledgments

We thank Xiao Liu for providing *Neurospora crassa* strains. The work was supported by the China National Key Research and Development Program (Grant no. 2024YFC3406000), the Chinese Academy of Sciences (CAS) Projects for Young Scientists in Basic Research (Grant no. YSBR-080), the Strategic Priority Research Program of Chinese Academy of Sciences (Grant no. XDA0450000) to C-M.S., and the Young Scientists Fund of the National Natural Science Foundation of China (Grant no. 32402338) to H.W. The funder has not played any role in the study design, data collection and analysis, the decision to publish, or the preparation of the manuscript.

## Author Contributions

**H.W.**: methodology, investigation, data curation, formal analysis, writing-original draft, and writing—review and editing;

**Y.T.**: methodology, investigation, data curation, formal analysis, writing-original draft, and writing—review and editing;

**J.M.**: biostatistics analysis, data curation, writing-original draft, and writing—review and editing;

**J.Y.**: formal analysis, writing—original draft, and writing-review and editing;

**M.L.**: formal analysis, writing—original draft, and writing-review and editing;

**P.J.**: biostatistics analysis, writing-original draft, and writing—review and editing;

**S.L.**: supervision, investigation, resources, data curation;

**H.X.**: methodology, investigation, data curation, finish the experiments involving *Fusarium graminearum*;

**G.T.**: methodology, investigation, data curation, finish the experiments involving *Fusarium graminearum*;

**W.L.**: supervision, methodology, data curation, finish the experiments involving *Fusarium graminearum*;

**H-S.G.**: supervision, project administration, methodology, resources, data curation, writing-review and editing.

**C-M.S.**: conceptualization, supervision, project administration, methodology, resources, data curation, writing-original draft, writing-review and editing.

All authors have read and agreed to the published version of the manuscript.

## Data Availability

All ChIP-Seq data were obtained from the SRA database (Home - SRA - NCBI). The ChIP-Seq data for H3K9me3 modification in *V. dahliae* were downloaded from SRR10571946 and SRR10571947, respectively, and data for H3K27me3 modification were downloaded from SRR10571948 and SRR10571949. For H3K9me3 modification in *N. crassa*, the ChIP-Seq data were downloaded from SRR1566112 and SRR12229310. The ChIP-Seq data for H3K4me3 modification in *F. graminearum* were downloaded from SRR21677668 and SRR21677669. The reference genomes used in the mapping process are as follows: PRJNA1212841 for *V. dahliae* V592, GCF_000182925.2 for *N. crassa*, FgraminearumPH-1 for *F. graminearum* (available at FungiDB), and GCA_001010845.1 for *S. scitamineum*. The fCUT&Tag-Seq data and ChIP-Seq data used in this study are available in the NCBI BioProject database under accession numbers PRJNA1213569, PRJNA1166993, and PRJNA1248591.

## Ethics statement

No animal experiments nor human samples were involved in this study.

## Conflicts of Interest

The authors declare that there is no conflict of interest.

## Notes

https://dataview.ncbi.nlm.nih.gov/object/PRJNA1213569

### Competing Interest Statement

The authors have declared no competing interest.

### Summary of Updates

Accepted manuscript. The main text and Figures have been revised.

