## Supplementary_Figures for "fCUT&Tag-Seq: An Optimized CUT&Tag-based Method for High-Resolution Profiling of Histone Modifications and Chromatin-Binding Proteins in Fungi"

A

| Sample | SeqNum | SumBase | N50Len | N90Len | MeanLen | MaxLen |
|---|---|---|---|---|---|---|
| *Verticillium dahliae* V592 | 156947206 | 40711754401 | 452 | 136 | 64.849 | 9518 |

B

| Chr | Chr1 | Chr2 | Chr3 | Chr4 | Chr5 | Chr6 | Chr7 | Chr8 |
|---|---|---|---|---|---|---|---|---|
| Length | 4416504 | 7834363 | 5191406 | 4182892 | 3710032 | 3337687 | 3268461 | 3754928 |
| GC% | 52.88 | 54.63 | 51.5 | 52.89 | 53.32 | 53.11 | 52.31 | 55.27 |

C

| Sample | Gene Number | CDS Averange length | CDS Averange GC(%) |
|---|---|---|---|
| *Verticillium dahliae* V592 | 8332 | 1435.8863 | 59.55 |

D

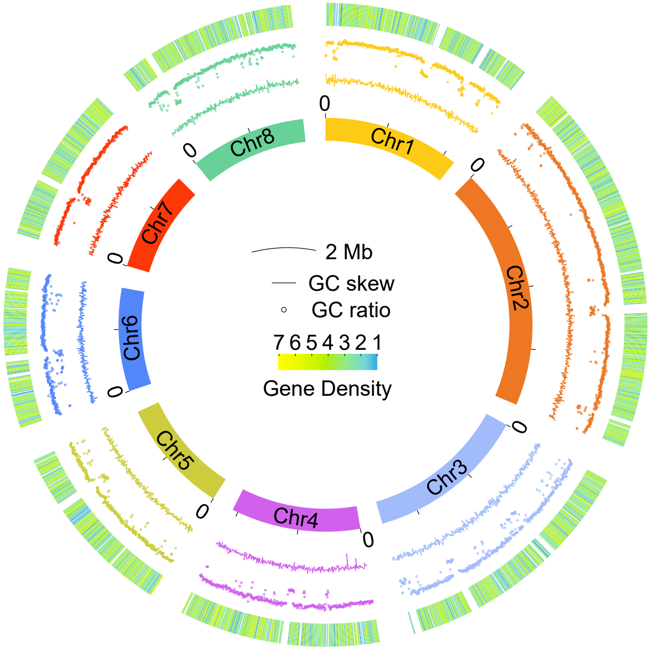

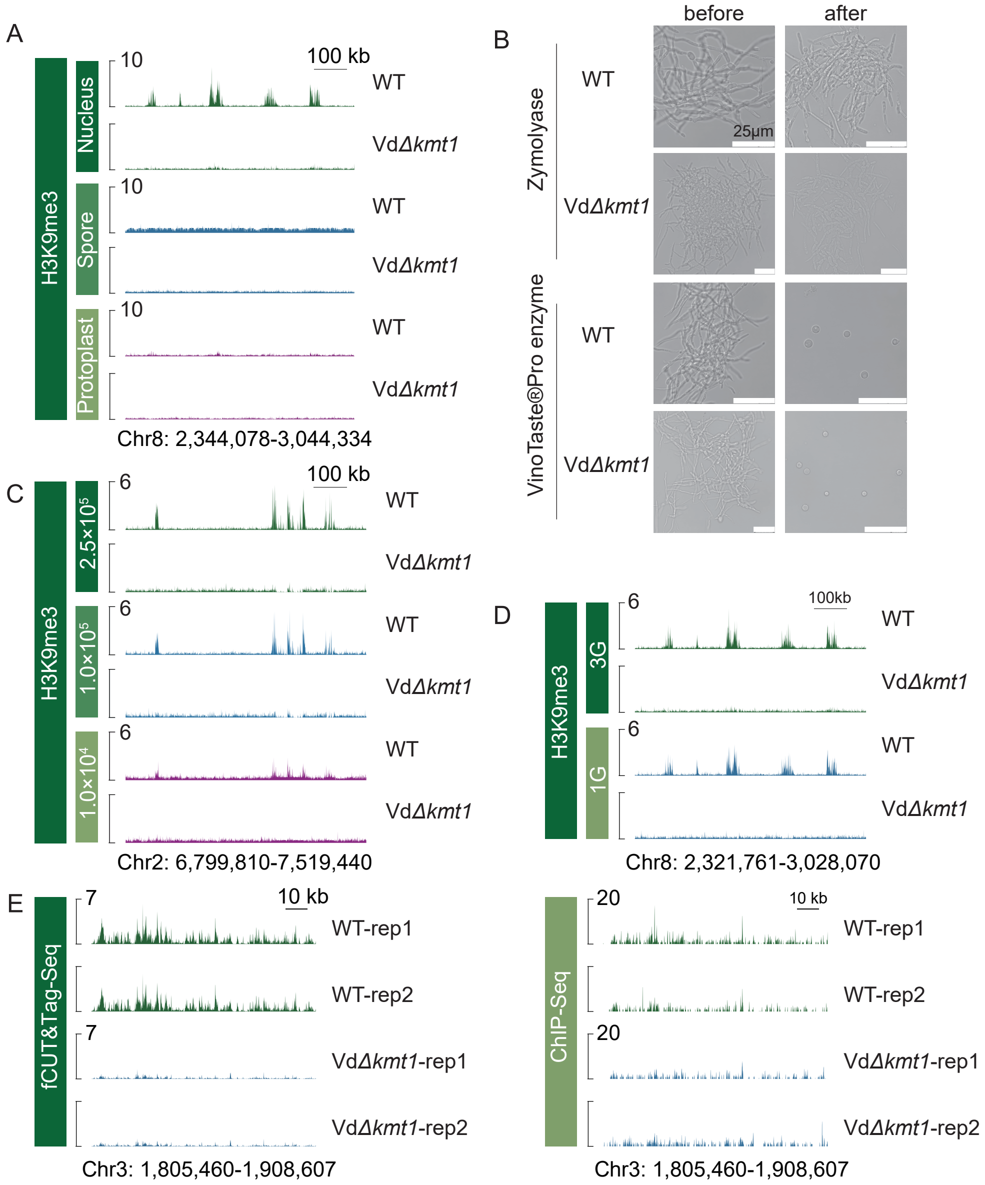

A
H3K9me3
Nucleus
Spore
Protoplast
10
100 kb
WT
VdΔkmt1
Chr8: 2,344,078-3,044,334
B
before
after
Zymolyase
VinoTaste®Pro enzyme
WT
VdΔkmt1
25μm
C
H3K9me3
2.5×10⁵
1.0×10⁵
1.0×10⁴
6
100 kb
WT
VdΔkmt1
Chr2: 6,799,810-7,519,440
D
H3K9me3
3G
1G
6
100kb
WT
VdΔkmt1
Chr8: 2,321,761-3,028,070
E
fCUT&Tag-Seq
7
10 kb
WT-rep1
WT-rep2
VdΔkmt1-rep1
VdΔkmt1-rep2
Chr3: 1,805,460-1,908,607
ChIP-Seq
20
10 kb
WT-rep1
WT-rep2
VdΔkmt1-rep1
VdΔkmt1-rep2
Chr3: 1,805,460-1,908,607

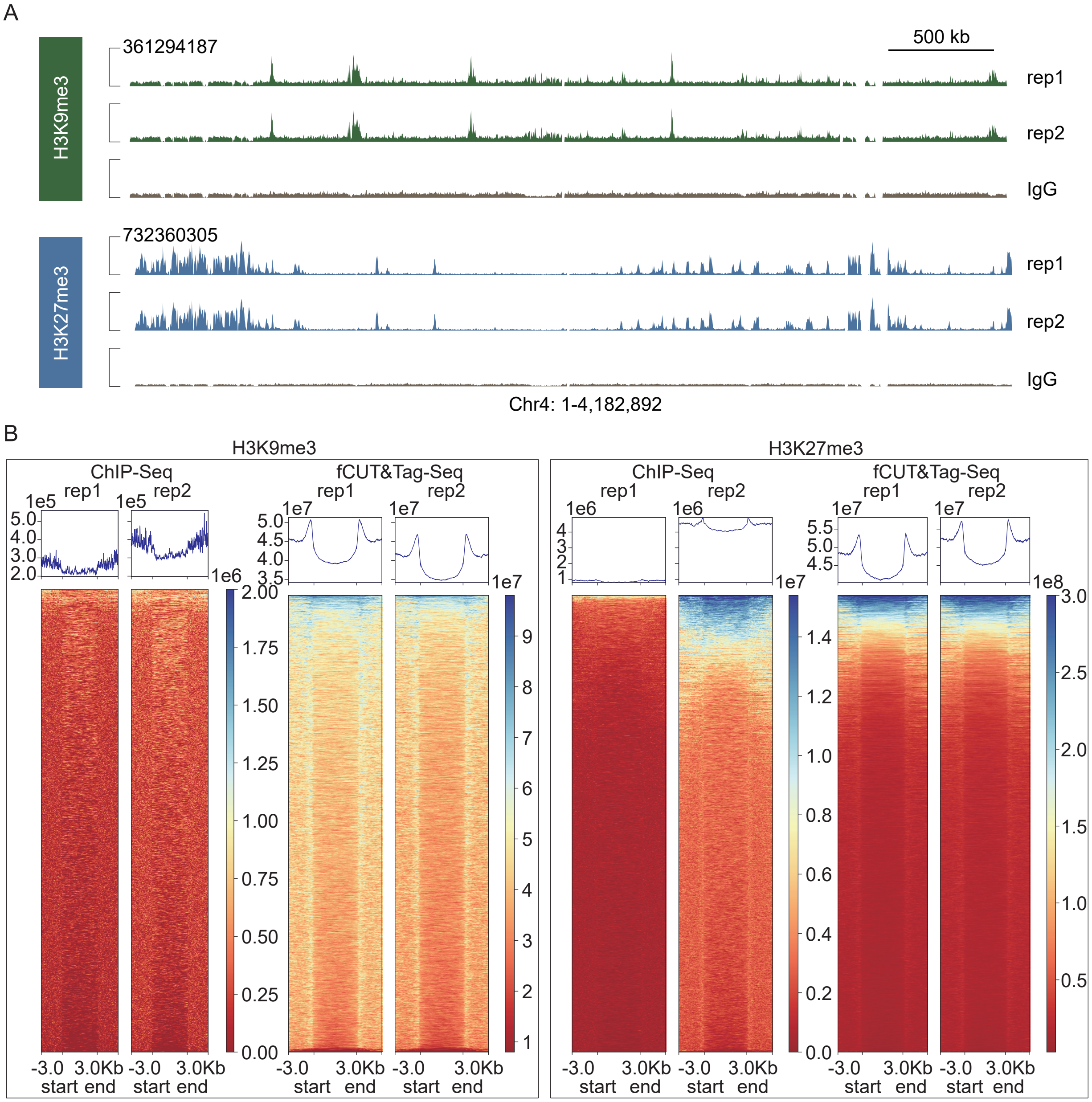
A
500 kb
H3K9me3
361294187
rep1
rep2
IgG
H3K27me3
732360305
rep1
rep2
IgG
Chr4: 1-4,182,892
B
H3K9me3
ChIP-Seq
fCUT&Tag-Seq
H3K27me3
ChIP-Seq
fCUT&Tag-Seq
rep1
rep2
-3.0
3.0Kb
start
end

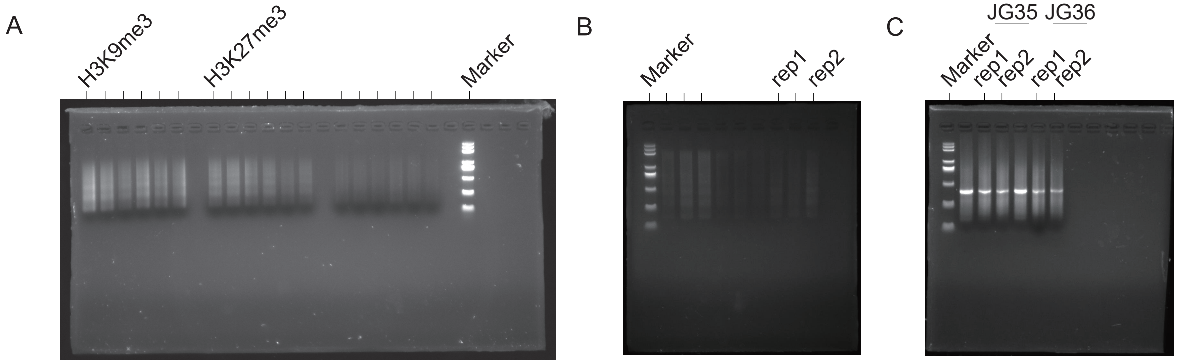
A
H3K9me3
H3K27me3
Marker
B
Marker
rep1
rep2
C
JG35
JG36
Marker
rep1
rep2
rep1
rep2

**Table S1: Primers used in this study.**

| Primers | Sequences (5'→3') | Application |
|---|---|---|
| VdKMT1-A | AATTCGAGCTCGCTGAGGGTTTAATTAACAACATTACGCAGCTCTACGAGT | Vd*Δkmt1* construct |
| VdKMT1-a | TCGATGGGCCCGCTGAGGACTTAATTAAGTTGACTGATCTGTAAATAAG | |
| VdKMT1-d | CCCCGACTAGTGCTGAGGCATTAATTAACGAGGCACCGGATGGGAGGG | |
| VdKMT1-B | TCACGAAGCTTGCTGAGGTCTTAATTAAGTCGTGATGGTGCCTGTGTGGAG | |
| Hpt-F | TCTCCTTGCATGCACCATTCCTTG | Detection of Vd*Δkmt1* |
| HPT-R | GCAGCTATTTACCCGCAGGA | |
| VdKMT1-V1 | CTCAGGGCAACATTGGAGGT | |
| VdKMT1-V2 | CAAGACAGCAAGCACGAGAAG | |
| VdEZH2-A | TTCGAGCTCGCTGAGGGTTTAATTAAATTCAAGGGTTTGGATCGG | Vd*Δezh2* construct |
| VdEZH2-a | GATGGGCCCGCTGAGGACTTAATTAATGCGAATCTGCTTGTGAGG | |
| VdEZH2-d | CCGACTAGTGCTGAGGCATTAATTAAGGATTTCCTGCCATGCAC | |
| VdEZH2-B | ACGAAGCTTGCTGAGGTCTTAATTAAAGGGTAAGGCAGGGACCG | |
| Hpt-F | TCTCCTTGCATGCACCATTCCTTG | Detection of Vd*Δezh2* |
| HPT-R | GCAGCTATTTACCCGCAGGA | |
| VdEZH2-V1 | GGCATGGTGTGTAATTGAAA | |
| VdEZH2-V2 | TGTGACCTCGCCGCTATTG | |
| SsGcn5-tg-A | AGATGACCGTGAACCACTCG | SsGcn5-Flag construct |
| SsGcn5-tg-a | CCGTCATGGTCTTTGTAGTCCAAGGAACTCTGCACCTTCG | |
| SsGcn5-d | GTCCGCAATGTGTTATTAAGTGGTCCCAAGGTTGTCTTG | |
| SsGcn5-B | GGTGTAGATGGCGAGGAGAT | |
| HPT-LB-F | GACTACAAAGACCATGACGG | |
| HPT-LB-R | GGTCAAGACCAATGCGGAGC | |
| HPT-RB-F | GCAAGACCTGCCTGAAACCG | |
| HPT-RB-R | CTTAATAACACATTGCGGAC | |
| SsGcn5-C (V1) | TCCGGGCCACACTCTTC | Detection of SsGcn5-Flag |
| SsGcn5-D (V2) | ATGGCAAGGTTGAGGCTGT | |
